# Improved nuclease-based prime editing by DNA repair modulation and pegRNA engineering

**DOI:** 10.1101/2024.02.01.578377

**Authors:** Panagiotis Antoniou, Louis Dacquay, Niklas Selfjord, Katja Madeyski-Bengtson, Anna-Lena Loyd, Euan Gordon, George Thom, Pei-Pei Hsieh, Sandra Wimberger, Saša Šviković, Mike Firth, Nina Akrap, Marcello Maresca, Martin Peterka

## Abstract

Prime editing is a genome engineering tool that allows installation of small edits with high precision. However, prime editing efficiency and purity can vary widely across different edits, genomic targets, and cell types. Prime editing nuclease (PEn) utilizes a fully active Cas9 instead of the nickase employed in conventional prime editors. PEn is capable of editing sites resistant to nickase-based prime editors but induces more undesired editing events. In this work, we introduce two strategies to enhance PEn precision and efficiency. First, we apply a small molecule approach, selectively modulating DNA repair pathways, to improve PEn precision up to 9.8-fold and reduce off-target editing by 90%. Second, through pegRNA engineering, we devise a strategy that mitigates unintended pegRNA scaffold integration, which is a common prime editing by-product, enhancing precision up to 3.5-fold. We apply this approach to a specific type of PEn editing mediated through non-homologous end joining and use it to achieve efficient and precise prime editing in multiple human cell lines, primary human hepatocytes, and mouse embryos. Together, this work presents two general strategies to improve prime editing, overcomes the limitations of current PEn editors, and provides reliable and precise genome editing outcomes, a pivotal requirement for therapeutic applications.

## Introduction

Prime editing is a next generation genome engineering approach that allows DNA donor-free installation of various types of small genomic edits including insertions, deletions, and substitutions (Chen and Liu, 2023). Compared to more traditional approaches for precise genome editing such as homology-directed repair (HDR), prime editing induces comparably lower levels of undesired editing outcomes and off-targets (Anzalone et al., 2019, Liang et al., 2023). A typical prime editor (PE) comprises a Cas9 nickase fused to a reverse transcriptase (RT) via a flexible linker (Anzalone et al., 2019). PE operates by reverse-transcribing an RNA template that is part of a prime editing guide RNA (pegRNA). In doing so, PE introduces a 3’ genomic flap, which contains a desired edit as well as a homology region which anneals and integrates into the targeted genomic locus. Prime editing is a highly modular method and numerous PE systems were developed for various genome editing applications (Chen and Liu, 2023). Currently, broad adaptation of PE is impeded by several limitations, such as the need for extensive pegRNA optimizations and the widely varying editing efficiency observed across different types of edits, target sites, and cell types (Anzalone et al., 2019, Chen et al., 2021, Koeppel et al., 2023, Yu et al., 2023).

Recently, several studies reported a PE variant that is based on the fusion of RT to the wild-type, fully active *Sp*Cas9 nuclease (termed Prime Editing nuclease – PEn) (Peterka et al., 2022, Adikusuma et al., 2021, Kweon et al., 2023, Li et al., 2023, Jiang et al., 2021, Tao et al., 2022). PEn efficiently rescues editing at target sites resistant to nicking-based PE, likely due to a different and more robust DNA repair mechanism promoted by the presence of DNA double strand breaks (DSBs). However, DSB repair also leads to increased insertion and deletion (indel) formation and imprecise prime edits caused by the activity of end-joining pathways such as non-homologous end joining (NHEJ) and alternative end-joining (alt-EJ). In addition to PEn editing with pegRNAs, we previously introduced an alternative approach termed primed insertions (PRINS), which combines PEn and a special pegRNAs design (single PRINS guide RNA or springRNA). springRNA omits the homology region in the RT template required for flap annealing and efficiently promotes targeted insertions directly at the DSBs via the broadly active and highly efficient NHEJ pathway. While PRINS could be particularly relevant in post-mitotic tissues, which are typically deficient in homologous recombination but retain high NHEJ activity (Zhao et al., 2020, Yeh et al., 2019), it suffers from increased imprecise prime edits caused by RT readthrough into the single guide (sgRNA) scaffold, causing the final intended edit to contain additional sgRNA scaffold nucleotides (Peterka et al., 2022). Variable levels of scaffold integration have also been observed for nickase-based PEs (Anzalone et al., 2019, Petri et al., 2021, Fiumara et al., 2023). Thus, while PEn can be more robust and efficient than nicking PE, the induction of DSBs generally leads to increased levels of both indels and imprecise prime edits, which can limit the scope of PEn editing.

Here we present two methods that improve PEn editing by addressing limitations of the original PEn system. First, to reduce by-products of PEn editing with pegRNAs, we introduce ‘2 inhibitor’ PEn (2iPEn), a small molecule approach that uses a combination of inhibitors of DNA repair pathways, modulating the choice of DSB repair during PEn editing. We and others have previously demonstrated the advantages of employing dual NHEJ/alt-EJ inhibition in HDR-based methods to enhance editing efficiency and precision while simultaneously reducing off-target editing (Wimberger et al., 2023, Schimmel et al., 2023, Riesenberg et al., 2023, Arai and Nakao, 2021). The small molecule inhibitors target the core NHEJ and alt-EJ factors DNA-dependent protein-kinase (DNA-PK) and DNA polymerase Θ (PolΘ), respectively, thereby minimizing undesired indels formed by these pathways. Indeed, while investigating the mechanism of PEn editing, we have previously shown that knocking out PolΘ did not diminish PEn editing but further reduced indels observed upon DNA-PK inhibition (Peterka et al., 2022). With 2iPEn we exploit this principle to substantially improve the precision of PEn editing outcomes and decrease off-target editing, while maintaining high on-target efficiency. Our second strategy focuses on enhancing a special type of NHEJ-mediated PEn editing and involves a modified pegRNA design that includes an abasic spacer between the RT template and the sgRNA scaffold. This modification prevents the incorporation of unwanted scaffold sequences by blocking RT readthrough into the sgRNA scaffold and significantly improves editing precision and efficiency across multiple systems including primary human hepatocytes and mouse embryos.

## Results

### Small molecule inhibitors of DNA repair improve nuclease prime editing

We previously demonstrated that treatment with the highly selective and potent DNA-PK inhibitor AZD7648 (Fok et al., 2019) (referred to as DNA-PKi) increases precision (proportion of precise prime edits compared to editing by-products) of PEn editing (Peterka et al., 2022). Similarly, Li *et al*. co-expressed PEn with a ubiquitin variant that inhibits tumour suppressor p53-binding protein 1 (53BP1) (uPEn) (Li et al., 2023). Both systems suppress NHEJ-mediated indels and imprecise prime edits, resulting in a similar increase in PEn editing precision and efficiency. Nevertheless, in both cases, the levels of by-products remain high, likely due to a shift to alt-EJ DNA repair and/or incomplete NHEJ inhibition. Based on these observations and recent work that utilized inhibitors of DNA-PK and PolΘ to improve HDR knock-ins (Wimberger et al., 2023, Schimmel et al., 2023), we hypothesized that simultaneous inhibition of DNA-PK and PolΘ with small molecules could further improve the precision of PEn editing. Thus, in addition to DNA-PKi for efficient NHEJ repression, we used PolΘ inhibitors PolQi1 and PolQi2 to suppress alt-EJ repair. PolΘ contains a helicase-like domain at its N-terminus and a polymerase domain at its C-terminus, both of which are required for DSB repair (Wood and Doublié, 2022). PolQi1 targets the polymerase domain whereas PolQi2 inhibits the helicase domain of PolΘ (Wimberger et al., 2023). To test the effects of co-inhibition of DNA-PK and PolΘ on the precision of PEn editing, we installed point mutations and various insertions in HEK293T and HeLa cells using a panel of pegRNAs that were previously shown to be highly active but imprecise with the nuclease-based PEn or uPEn (Peterka et al., 2022, Li et al., 2023). Cells were edited with PEn in the presence of DNA-PKi alone, DNA-PKi in combination with PolQi1, or DNA-PKi in combination with PolQi1 and PolQi2. We compared editing outcomes for each of these conditions with uPEn (PEn-P2A-i53 configuration) (Li et al., 2023) as well as nickase-based PE5, which involves expression of dominant negative MLH1 suppressing DNA mismatch repair and an additional nicking sgRNA (Chen et al., 2021). As reported previously, while PEn and uPEn allowed efficient editing with pegRNAs that did not perform well with PE5, both suffered from increased levels of indels and imprecise prime edits (Figure 1a and 1b, Supplementary Figure 1a). As we have shown previously, PEn editing in the presence of DNA-PKi (“1iPEn”) increased the precision compared to PEn alone (Figure 1c) but did not mitigate indels unrelated to prime edits (Figure 1a, b), which were shifted from insertions towards deletions, suggesting a shift from NHEJ to alt-EJ-mediated DNA repair (Supplementary Figure 1a). Co-administration of DNA-PKi with PolQi1 (“2iPEn”) further improved editing precision compared to uPEn, PEn and 1iPEn (Figure 1c) by reducing indels and imprecise prime edits (Figure 1a and b, Supp 1a). Importantly, this effect was consistent across different target sites and cell lines (Figure 1c). Next, to inhibit both the polymerase and helicase-like domains of PolΘ, we combined DNA-PKi with PolQi1 and PolQi2 (“2^+^iPEn”). Remarkably, 2^+^iPEn consistently led to near complete editing purity, reaching the precision levels of PE5 (Figure 1c), while maintaining the high efficiency of PEn (Figure 1a and b). The precision increase of 2^+^iPEn compared to PEn and uPEn was 4.3-fold and 2.1-fold respectively in HEK293T cells and 9.8-fold and 7.1-fold in HeLa cells (Figure 1c). In summary (Figure 1d), we demonstrate how combining small molecule inhibitors to target both NHEJ and alt-EJ significantly improves the editing outcome of PEn by reducing by-products to the level of nickase-based PEs while maintaining the robustness and editing efficiency of PEn.

**Figure 1:**
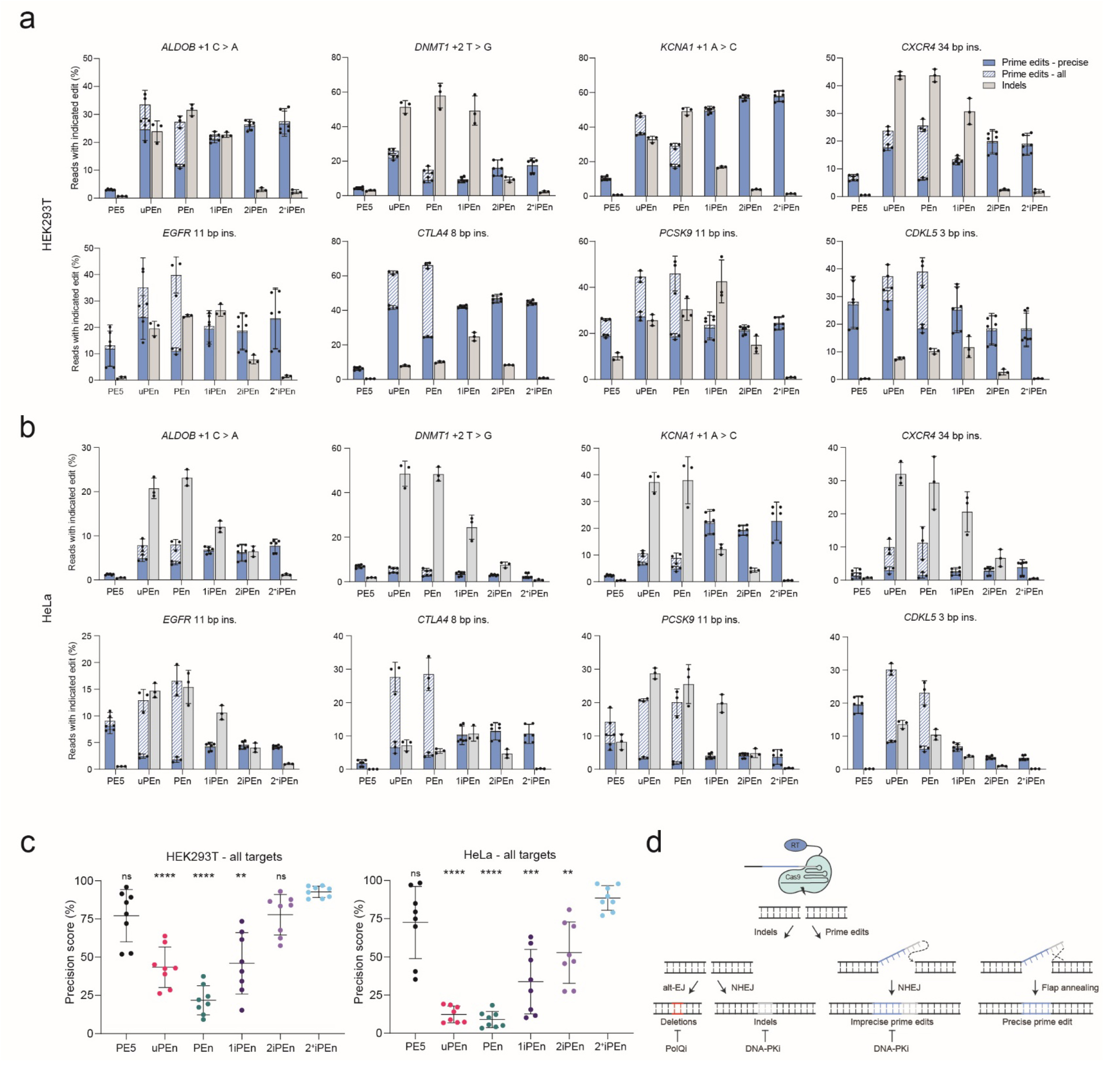
Co-inhibition of DNA-PK and PolΘ increases precision of PEn editing. Genome editing with six different prime editing systems along with eight pegRNAs installing point mutations or insertions (ins) using plasmid co-transfection in **a)** HEK293T and **b)** HeLa cells. Editing outcomes were analyzed by targeted amplicon sequencing and quantified using CRISPResso2. Plots show mean ± SD *of n* = 3 biological replicates. “Prime edits – all” and “Prime edits – precise” categories are superimposed. **c)** Precision of six different prime editing systems in HEK293T and HeLa cells across all loci tested in panels 1a and 1b. “Precision score” was calculated as the total number of NGS reads with precise prime edit per overall editing. Error bars represent mean ± SD. *P*-values were determined using one-way ANOVA (two-tailed, paired). Displayed pairwise comparisons are relative to 2^+^iPEn. Calculated *P* values HEK293T cells 2^+^iPEn vs. PE5 = 0.1564; 2^+^iPEn vs uPEn <0.0001; 2^+^iPEn vs. PEn <0.0001; 2^+^iPEn vs. 1iPEn = 0.001; 2^+^iPEn vs. 2iPEn = 0.0991. Calculated *P* values Hela cells 2^+^iPEn vs. PE5 = 0.4296; 2^+^iPEn vs uPEn <0.0001; 2^+^iPEn vs. PEn <0.0001; 2^+^iPEn vs. 1iPEn = 0.0002; 2^+^iPEn vs. 2iPEn = 0.0017. *P* values for all other comparisons are provided as Supplementary Data 1. **d)** Schematic representation of 2iPEn depicting molecular mechanisms of four possible PEn editing outcomes and their modulation by DNA repair inhibitors. RT = reverse transcriptase.

### Simultaneous DNA-PK and PolΘ inhibition reduces PEn off-target editing

We have previously shown that PEn can cause increased editing at individual off-target sites using promiscuous pegRNAs compared to editing with Cas9 alone (Peterka et al., 2022) and a similar trend has been observed for uPEn (Li et al., 2023). This was at least in part due to the promiscuous priming by RT at the off-target cut leading to its inability to be seamlessly repaired back to the original wild-type state (Peterka et al., 2022). As we have shown earlier that co-inhibition of DNA-PK and PolΘ substantially mitigates off-target editing in Cas9-treated samples (Wimberger et al., 2023), we wondered whether editing with 1iPEn, 2iPEn or 2^+^iPEn will result in a similar alleviating effect on PEn off-target editing. To explore the impact of small molecule inhibitors of DNA repair on PEn off-target editing, we systematically investigated four promiscuous pegRNAs targeting the *EMX1, FANCF, HEK3*, and *HEK4* loci together with 13 corresponding off-target sites (Anzalone et al., 2019, Tsai et al., 2015) using targeted amplicon sequencing (Figure 2a). Consistent with our prior findings, PEn editing showed cleavage activity across all interrogated off-target sites. Inhibition of 53BP1 (uPEn) or DNA-PK (1iPEn) equally reduced off-target editing at all tested sites by about 35-40% on average. Notably, the incorporation of PolΘ inhibitor (2iPEn) further diminished off-target editing by about 74% on average compared to PEn editing alone. The most beneficial off-target profile was achieved when utilizing the combination of DNA-PKi and both PolΘ inhibitors (2^+^iPEn) leading to an average of 90% reduction in off-target editing. A complete overview of all on- and off-target editing events is provided in Supplementary Figure 2. To account for variable on-target editing levels across different PEs, we computed specificity scores to normalize off-target editing to on-target editing levels. The specificity score is calculated as 1 minus the ratio of editing frequency at the off-target site to that at the on-target site resulting in a range between 0 and 1. A specificity score of 1 denotes the absence of off-target editing (Kim et al., 2023). This analysis substantiated the off-target editing results, revealing a significant decrease in off-target editing for 2iPEn and 2^+^iPEn compared to PEn (Figure 2B). In summary, our findings demonstrate a substantial reduction of PEn off-target editing through the concurrent inhibition of DNA-PK and PolΘ.

**Figure 2:**
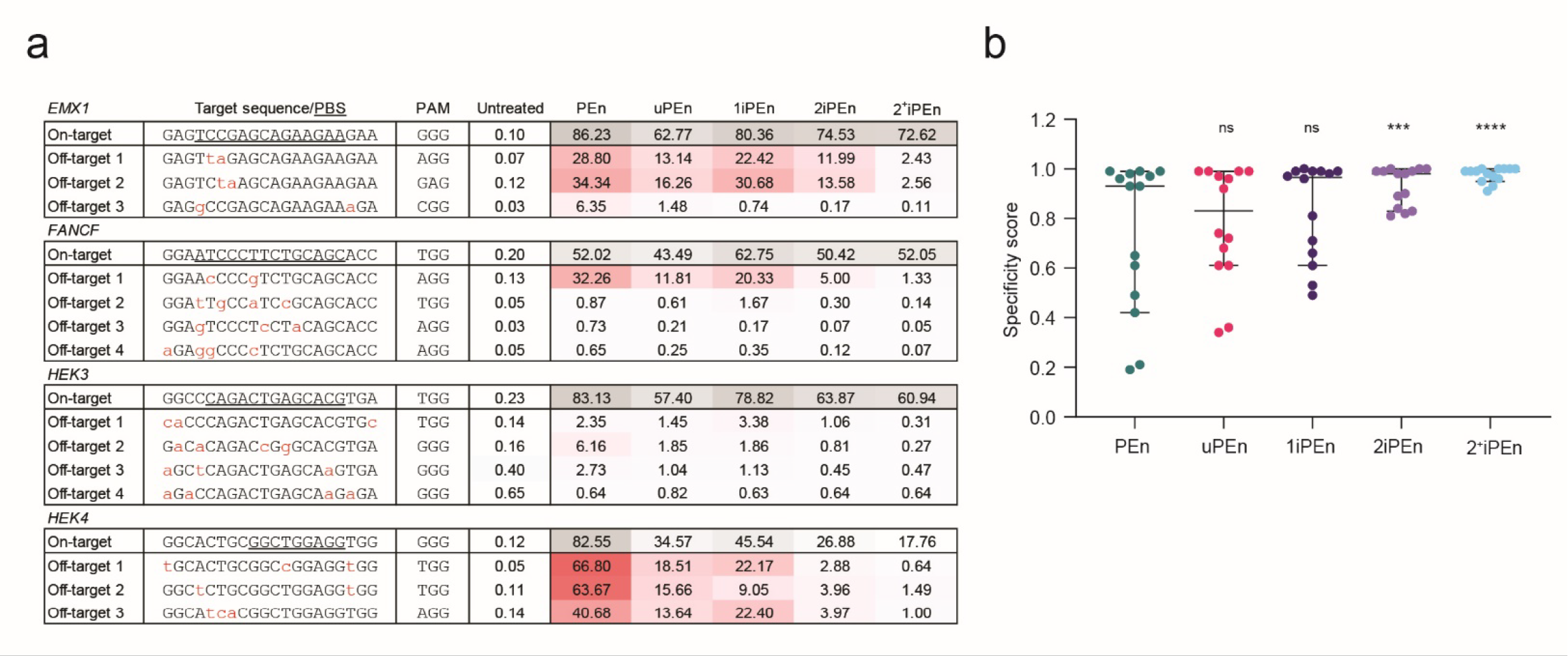
Co-inhibition of DNA-PK and PolΘ decreases PEn off-target editing. **a)** Targeted amplicon sequencing and CRISPResso 2 analysis at four on-target and 13 off-target sites for the indicated PEn configurations using plasmid co-transfections in HEK293T cells. Editing is shown as percentages of modified reads in each sample. Mismatches to the on-target gRNA sequence are highlighted in red, and the PBS region is underlined. **b)** Computed specificity scores for all off-target data shown in 2a). Each dot in the graph represents one individual off-target, horizontal lines represent median and 95% confidence intervals of *n* = 3 biological replicates with *n* = 3 technical replicates each. Significance levels of specificity scores were calculated with the non-parametric Friedman test (paired, two-tailed). Multiple comparisons were corrected using Dunn’s test. Calculated *P* values: PEn vs. uPEn >0.9999 (ns), PEn vs. 1iPEn = 0.7544 (ns), PEn vs. 2iPEn = 0.0008 (^***^), PEn vs. 2^+^iPEn <0.0001 (^****^). ns; non-significant.

### Mitigating pegRNA scaffold integrations by abasic spacer

Unwanted insertions of the pegRNA scaffold are one of the common prime editing by-products. After transcribing the RT template, RT normally terminates due to its collision with Cas9 and/or the first stem loop of the pegRNA scaffold. However, in some cases, reverse transcription can extend beyond the intended template, potentially incorporating variable lengths of pegRNA scaffold sequence. This may subsequently affect the efficiency and precision of prime editing. The unwanted scaffold integrations were observed in the original PE work and since then have been found to be variably pronounced in different prime editing contexts (Anzalone et al., 2019, Fiumara et al., 2023, Petri et al., 2021). Here we focus on PRINS prime editing, which employs a robust NHEJ-based mechanism for programmed insertions. However, because of its specific homology-free pegRNA design (springRNA), PRINS is especially prone to unwanted scaffold insertions (Peterka et al., 2022). We hypothesized that a modified nucleotide in the springRNA, acting as a reverse transcription roadblock, would terminate RT at a defined position and prevent unwanted scaffold insertions. To this end, we developed spaceRNA – a modified springRNA that contains a single riboabasic spacer (rSpacer). This oligonucleotide modification is well known to block extension by various DNA polymerases (Boiteux and Guillet, 2004). We positioned the rSpacer between the intended insert in the RT template and the springRNA scaffold (Figure 3a). To test the effect of spaceRNA on PRINS editing precision and efficiency in different cell lines, we electroporated K562, HEK293T, HeLa, and HepG2 cells with PEn mRNA and synthetic springRNA or spaceRNA to install small insertions at *PCSK9* and *HBEGF* loci. At both targets, and in all cell lines tested, springRNA-mediated PEn editing led to high levels of overall prime editing, most of which contained additional nucleotides that matched the scaffold sequence (Figure 3b). The typical size of the incorporated scaffold was 1-3 nucleotides (Figure 3c). On the contrary, spaceRNA-mediated PEn editing showed a complete elimination of scaffold incorporation events, and significantly higher levels of precise prime edits measured by targeted amplicon sequencing (Figure 3b and d). The average precision across all four cell lines was 3.5-fold and 3.2-fold higher with spaceRNA vs. springRNA at *PCSK9* and *HBEGF* sites, respectively (Figure 3d). Taken together, these data demonstrate that an rSpacer positioned directly after the RT template prevents unwanted scaffold integrations and substantially increases the precision of PRINS.

**Figure 3:**
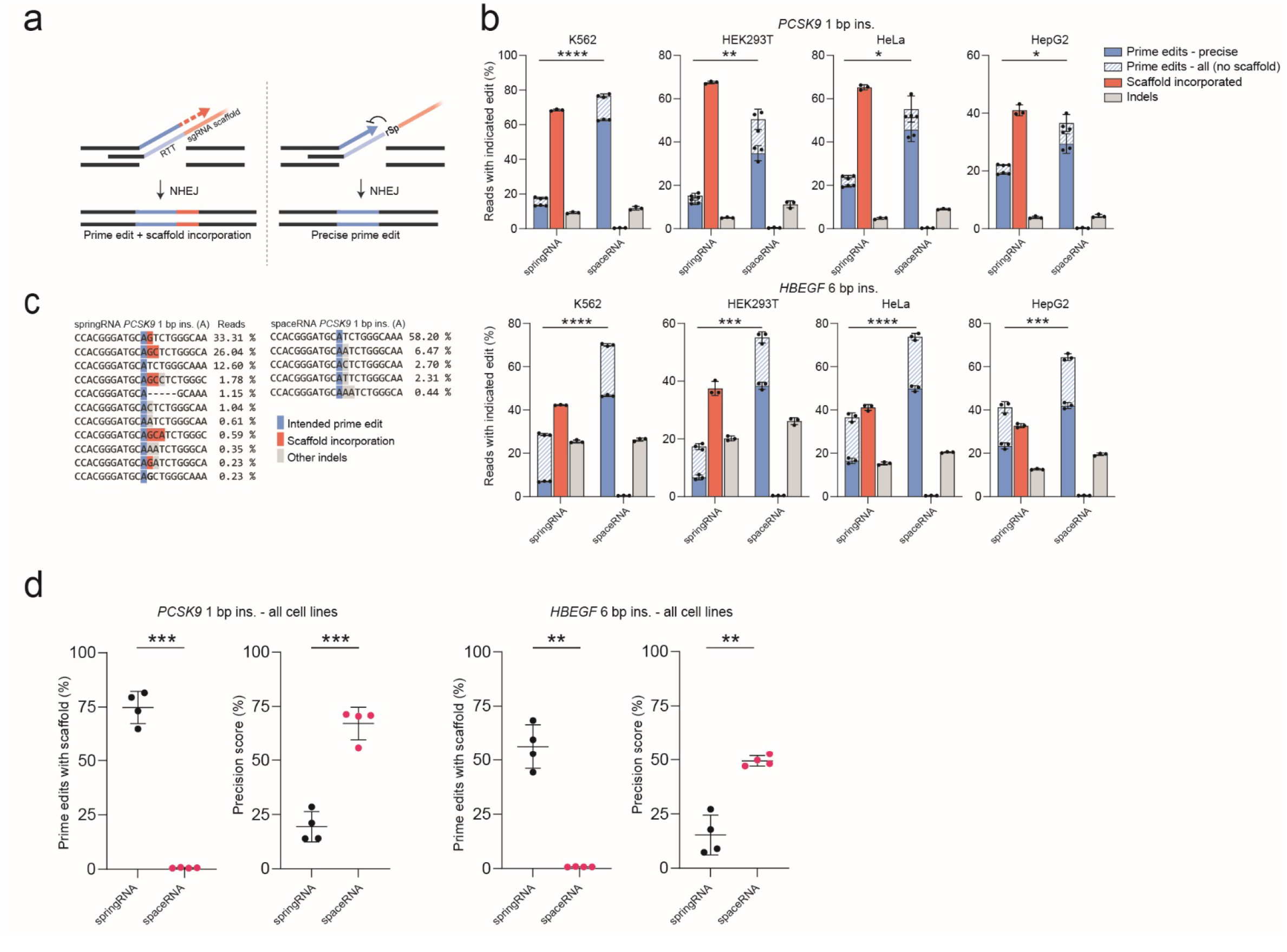
Elimination of pegRNA scaffold integrations through a riboabasic spacer. **a)** Schematic representation of PRINS editing. Left: regular PRINS editing resulting in unintended sgRNA scaffold integration due to flap extension beyond intended RT template (RTT). Right: PRINS editing using spaceRNA with a single riboabasic spacer (rSp) following RTT prevents polymerase readthrough into the sgRNA scaffold. **b)** PRINS editing at *PCSK9* and *HBEGF* genomic loci in four different human cell lines using electroporation of PEn mRNA in combination with synthetic springRNA or spaceRNA to install indicated small insertions (ins). Editing outcomes were analyzed by targeted amplicon sequencing and quantified using CRISPResso2. Plots show mean ± SD of *n* = 3 biological replicates. Statistical difference for “Prime edits – precise” was determined using Student’s t-test (paired, two-tailed). ^*^*P* < 0.05, ^**^*P* < 0.01, ^***^*P* < 0.001. Calculated *P* values: *PCSK9* (K562) < 0.0001, *PCSK9* (HEK293T) = 0.0061, *PCSK9* (HeLa) = 0.0137, *PCSK9* (HepG2) = 0.0345, *HBEGF* (K562) < 0.0001, *HBEGF* (HEK293T) = 0.0002, *HBEGF* (HeLa) < 0.0001, *HBEGF* (HepG2): 0.0003. **c)** Detailed breakdown of *PCSK9* editing in K562 cells from Fig. 3b. The alignment shows NGS reads containing prime edited alleles and their frequencies after editing with PEn and springRNA (top portion of the alignment) or spaceRNA (bottom portion of the alignment). **d)** Quantification of data from Fig 3b. “Prime edits with scaffold” were calculated as total number of NGS prime edited reads with scaffold integration per total prime edits. “Precision score” was calculated as the total number of NGS reads with precise prime edit per overall editing. Error bars represent mean ± SD. Statistical difference was determined using Student’s t-test (paired, two-tailed). ^*^*P* < 0.05, ^**^*P* < 0.01, ^***^*P* < 0.001. Calculated *P* values: *PCSK9* “Prime edits with scaffold” = 0.0003, *PCSK9* “Precision score” = 0.0008, *HBEGF* “Prime edits with scaffold” = 0.0017 *HBEGF* “Precision score” = 0.0034.

### PRINS editing in primary human hepatocytes

PRINS installs insertions via NHEJ, the predominant DSB repair pathway in most human cell types operating independently of cell cycle progression (Zhao et al., 2020). Hence, we hypothesized that PRINS could be a useful tool for precise editing of postmitotic cells. To test this, we applied PRINS for editing of primary human hepatocytes (PHHs). We transfected PHHs with PEn mRNA and a corresponding springRNA or spaceRNA to install small insertions at the *HBEGF* or *PCSK9* loci. Analogous to cell lines, targeted amplicon sequencing analysis revealed high overall prime editing with a substantial portion of additional scaffold sequences, which were mitigated using spaceRNA, leading to improved precision as well as efficiency, reaching up to 40% of precise editing at the therapeutically relevant *PCSK9* locus (Figure 4a).

**Figure 4:**
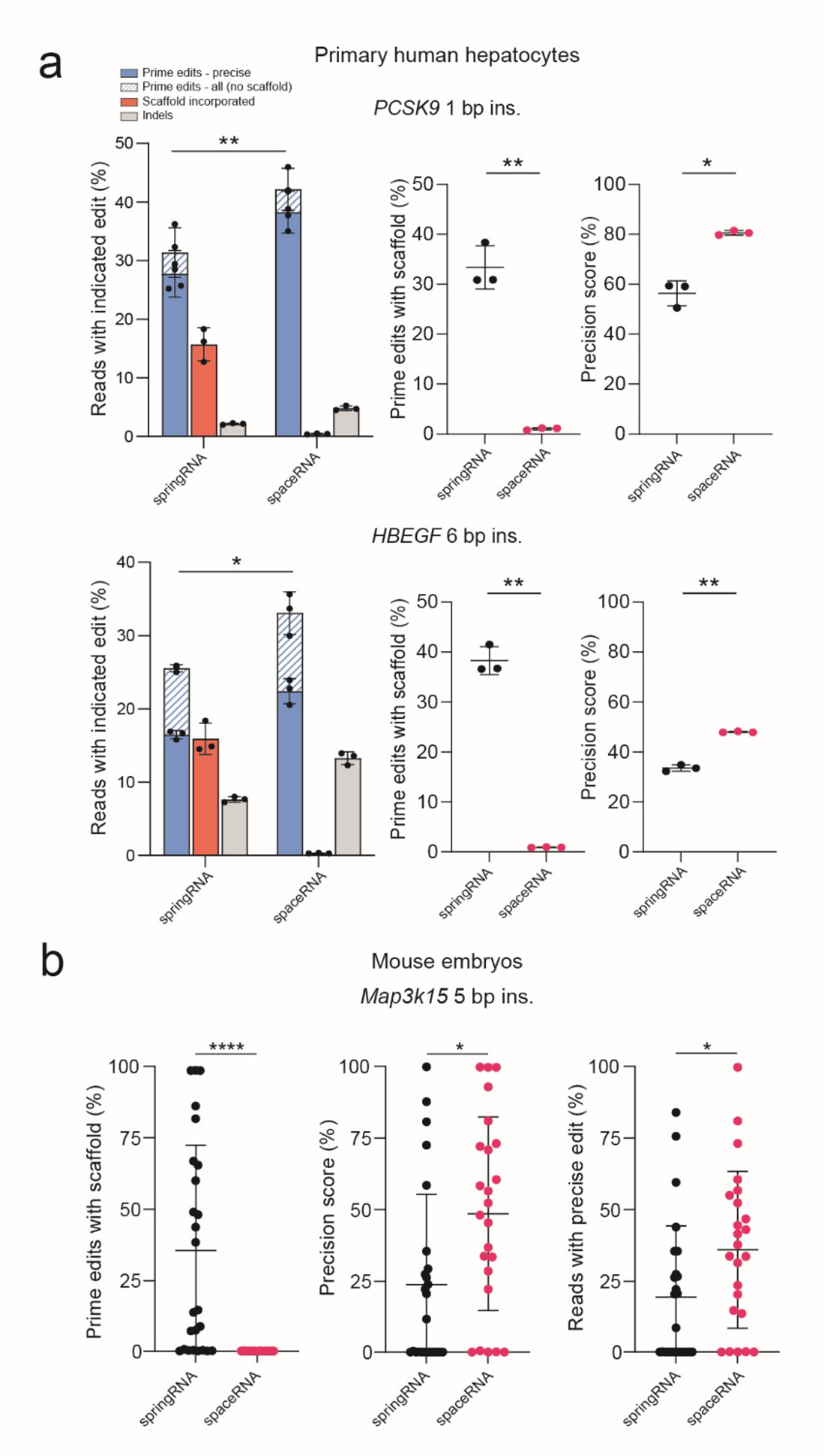
NHEJ-based PRINS editing efficiently installs small insertions in primary human hepatocytes and mouse embryos. **a)** PRINS editing at *PCSK9* and *HBEGF* genomic loci in primary human hepatocytes using transfection of PEn mRNA in combination with a synthetic springRNA or spaceRNA to install indicated small insertions (ins.). Editing outcomes were analyzed by targeted amplicon sequencing and quantified using CRISPResso2. “Prime edits with scaffold” were calculated as total number of NGS prime edited reads with scaffold integration per total prime edits. “Precision score” was calculated as the total number of NGS reads with precise prime edit per overall editing. Plots show mean ± SD of *n* = 3 biological replicates. Statistical difference was determined using Student’s t-test (paired, two-tailed). ^*^*P* < 0.05, ^**^*P* < 0.01, ^***^*P* < 0.001. Calculated *P* values: *PCSK9* “Prime edits - precise” = 0.005, *PCSK9* “Prime edits with scaffold” = 0.0057, *PCSK9* “Precision score” = 0.0104, *HBEGF* “Prime edits - precise” = 0.0123, *HBEGF* “Prime edits with scaffold” = 0.0018 *HBEGF* “Precision score” = 0.0027. **b)** PRINS editing of one-cell mouse embryos using electroporation of PEn-springRNA or PEn-spaceRNA RNP complexes to install a 5 -bp insertion into the endogenous *Map3k15* locus. Each data point represents a single embryo edited with either springRNA (n = 25) or spaceRNA (n = 24). Single embryos were analyzed after 5 days *in vitro* cultivation by targeted amplicon sequencing and the editing outcomes were quantified using CRISPResso2. “Prime edits with scaffold” were calculated as total number of NGS prime edited reads with scaffold integration per total prime edited reads. “Precision score” was calculated as the total number of NGS reads with precise prime edit per overall editing. “Reads with precise edit” corresponds to the total number of precisely edited NGS reads in each embryo. Error bars represent mean ± SD. Statistical significance was determined using Student’s t-test (unpaired, two-tailed). ^*^*P* < 0.05, ^**^*P* < 0.01, ^***^*P* < 0.001. Calculated *P* values: “Prime edits with scaffold” <0.0001, “Precision score” = 0.011, “Reads with precise edit” = 0.0319.

### PRINS editing in mouse embryos

NHEJ is highly active in mouse embryos (Singh et al., 2015). We thus speculated that PRINS might be an effective method for precise genome editing in this system. We electroporated single-cell mouse embryos with PEn-springRNA or PEn-spaceRNA ribonucleoprotein complexes to install a 5 bp insertion into the endogenous *Map3k15* locus. The electroporated embryos were cultured *in vitro* for 5 days after which they were lysed and subjected to targeted amplicon sequencing. The analysis of the *Map3k15* locus sequencing revealed variable precision and efficiency of PRINS editing in both conditions (Figure 4b and Supplementary figure 4). Notably, spaceRNA increased precision 2-fold and efficiency 1.8-fold compared to editing with unmodified springRNA. In the spaceRNA group, 7 out of 24 embryos analyzed contained precise prime edits in at least 50%, reaching up to 100% of sequenced reads. Taken together, these results serve as a basis for further exploration of PRINS as an efficient method to install small insertions in mouse embryos.

## Discussion

Prime editing has emerged as a groundbreaking technique for precisely repairing small genomic mutations and indels, making it a valuable tool in disease modeling and genetic disease treatment. Unlike HDR, which is constrained by the cell cycle, and base editing, which is limited in addressing specific point mutations, prime editing offers an alternative to address a broader range of genetic alterations. Yet, while it offers remarkable potential, particularly in non-dividing cells, there is a need to enhance its efficiency to fully harness its capabilities.

In this study, we present two general strategies to achieve highly efficient prime editing in cell lines, postmitotic primary cells, and mouse embryos exploiting both homology-dependent and homology-independent prime editing. We build upon nuclease prime editor PEn, which displays high efficiency at nickase-based PE resistant sites but lacks the precision of nickase-based PE. This is caused by indel formation and various types of imprecise prime edits arising from DSB repair.

First, we applied small molecules to inhibit NHEJ and alt-EJ pathways that give rise to undesired by-products of PEn editing. To this end, we adapted a small molecule cocktail previously reported to improve the outcomes of HDR editing. The NHEJ-inhibiting component of this cocktail is AZD7648 – a highly potent and selective DNA-PK inhibitor that targets the ATP-binding pocket of DNA-PK (Fok et al., 2019). For alt-EJ inhibition, we target PolΘ with two selective inhibitors – PolQi1 and PolQi2, which target the polymerase and the helicase domain of PolΘ, respectively (Wimberger et al., 2023). While DNA-PK inhibition alone can improve PEn editing by mitigating small indels and imprecise prime editing integrations, alt-EJ remains active and can compensate for NHEJ deficiency. The shift to alt-EJ repair leads to a change of indel composition, with a prevalence of deletions. Simultaneous inhibition of DNA-PK and PolΘ increased the precision of PEn up to 9.8-fold compared to editing with PEn alone and up to 7.1-fold compared to uPEn, reaching precision levels of nickase PE. Thus, using the 2iPEn strategy, we achieved near complete purity of precise prime edits while retaining high efficiency of PEn editing. Importantly, PEn was previously shown to display relatively high off-target editing. In this work, we demonstrated that small molecule inhibitor treatment significantly reduces PEn off-target editing by 90% on average. In summary, the 2iPEn approach improves the precision and safety of PEn and will be readily adaptable to other prime editing platforms.

Next, we focused on enhancing PRINS, a specific type of PEn editing which utilizes pegRNAs that lack a homology region in the RT template, termed springRNA. springRNAs install desired insertions directly at the Cas9 cut by precise NHEJ-mediated repair. The NHEJ insertion mechanism is an advantageous feature of PRINS, as this pathway has robust activity in different cell types, including postmitotic, non-cycling cells. However, the original version of PRINS frequently installs unwanted springRNA scaffold sequences due to the RT readthrough beyond the RT template. To address this issue, we developed spaceRNA – a springRNA design where the scaffold and RT template are separated by a riboabasic spacer acting as a transcriptional RT barrier. We show that spaceRNAs efficiently eliminate undesired scaffold incorporation, significantly enhancing the precision of PRINS editing up to 3.5-fold in different cell lines with precise editing efficiency reaching up to 60%. Importantly, we demonstrate that the improved PRINS system is highly effective in PHHs, which are fully differentiated postmitotic cells not amenable to modification through HDR and thus with limited options for precise genome editing. We use PRINS to achieve efficient and precise PHH editing at two loci including up to 40% of precise prime edits at the therapeutically relevant *PCSK9* locus. Lastly, we tested PRINS as a tool for editing mouse embryos, a system where Cas9-induced, NHEJ-mediated indels can be easily generated, but introduction of precise modifications using existing methods can be challenging. We show that PRINS efficiently installs small insertions in mouse embryos and is further improved by employing spaceRNAs. These proof-of-concept results show that PRINS has the potential to become a valuable tool for generation of genetically engineered mouse models.

While our current study demonstrates the successful application of the 2iPEn and spaceRNA strategies to PEn, it is important to note that these findings have broader implications. The challenges we have addressed, specifically mitigating indel formation and decreasing scaffold incorporation, are not unique to prime editing nucleases alone. Therefore, both 2i and spaceRNAs strategies are also applicable to enhance classical prime editing systems, potentially improving the overall efficiency and precision of this genome editing tool. In conclusion, this work expands the prime editing toolbox with two easily adaptable approaches that allow highly efficient and precise prime editing in cell lines, primary cells, and mouse embryos.

## Methods

### Ethical statement

AstraZeneca has a governance framework and processes in place to ensure that commercial sources have appropriate patient consent and ethical approval in place for collection of the samples for research purposes including use by for-profit companies.

### DNA and RNA constructs

PEmax (pCMV-PEmax-P2A-hMLH1dn (Chen et al., 2021), uPEn (uPEn3) (Li et al., 2023) and PEn were generated by gene synthesis (Genscript). The sequence of PEn corresponds to PEmax with the H840A mutation reversed to the original histidine, restoring nuclease activity. All pegRNAs and nicking sgRNAs were generated by gene synthesis and cloned in pMA vector together with an upstream U6 promoter (GeneArt). All pegRNA sequences used in this study are listed in Supplementary Data 1. Synthetic pegRNAs were synthesized by IDT. spaceRNAs with riboabasic spacer (rSpacer) were synthesized by Horizon discovery. Capped PEnmax mRNA was generated following T7 directed *in vitro* transcription using a linearized PEnmax DNA template. The *in vitro* transcription reaction produced fully modified mRNA, replacing uridines with N1-Methyl-pseudouridine and cap-1 capped mRNA using TriLink’s CleanCap-AG Cap1 analogue. The mRNA was subsequently column purified using MEGAClear transcription clean-up kit (ThermoFisher) and mRNA purity analyzed using a fragment analyzer (Agilent).

### Protein expression and purification

The sequence of PEn with a C-terminal his tag was cloned into pET24a. The expression plasmid was then transformed into *Escherichia coli* BL21lDE3 Star (Thermofisher) for use in protein production. Autoinduction protocol (Studier, 2005) was used for the over-production of PEn. Essentially, the culture was first grown over-night at 37 °C, before inoculation with 800 mL of ZYP autoinduction media, which was then grown at 37 °C with shaking until OD_600_ reached about 1-2. The temperature was then lowered to 18 °C and the culture was grown for a further 24 hours, before harvesting the cells by centrifugation. Cell pellets were stored at –80 °C until further use. The cell pellets were then resuspended in 20 mM HEPES, pH 7.5, 500 mM NaCl, 1 mM DTT, 10% glycerol, and lysed by one pass through an Emulsiflex C3 (Avestin). The lysate was clarified by centrifugation at 20,000 *g* for 20 minutes. The supernatant was supplemented with 10 mM imidazole and the lysate was loaded onto a 5 mL HiTrap column (Cytiva) equilibrated in the same buffer. The column was washed with 20 column volumes of 20 mM HEPES pH, 7.5, 500 mM NaCl, 1 mM DTT, 10% glycerol, 20 mM imidazole, before elution with 300 mM imidazole. The eluted protein was diluted to about 200 mM NaCl, before further purification on a 5 mL HiTrap Heparin SP column (Cytiva). Finally, the protein was further purified by size exclusion chromatography on a Superdex 200 (26/60) column (Cytiva) equilibrated in a buffer consisting of 20 mM HEPES, pH 7.5, 300 mM NaCl, 1 mM DTT, and 10% glycerol. The peak containing PEn protein was pooled, concentrated to 10 mg/mL, flash-frozen in liquid nitrogen and stored at -80 °C until required.

### Small molecule compounds and drug treatments

DNA-PK inhibitor AZD7648 was purchased from MedChemExpress (HY-111783). PolQi1 (WO2021028643) and PolQi2 (WO2020243459) were provided by AstraZeneca (Gothenburg, SE). All compounds were dissolved in dimethyl sulfoxide (DMSO) at a stock concentration of 10 mM.

### Cell culture

HEK293T (ATCC, CRL-3216) were maintained in DMEM-GlutaMAX with 10% fetal bovine serum (FBS) and 1% penicillin/streptomycin. HeLa cells (ATCC, CCL-2) were cultured in MEM-HEPES-GlutaMAX with 10% FBS and 1% penicillin/streptomycin. HepG2 (ATCC, HB-8065) were cultured in MEM-HEPES-GlutaMAX with 10% FBS, 1x non-essential amino acids, 1 mM sodium pyruvate and 1% penicillin/streptomycin. K562 cells (ATCC CRL-243) were maintained in RPMI-1640-GlutaMAX supplemented with 10% heat-inactivated FBS, 20 mM HEPES, 1 mM sodium pyruvate and 1% penicillin/streptomycin. All reagents were purchased from Gibco. Cells were maintained at 37 °C in a 5% CO_2_ atmosphere and regularly tested for mycoplasma contamination. Cell line identity was authenticated through STR profiling (IDEXX BioAnalytics). Primary human hepatocytes (LifeNet Health) were thawed 1 day before transfection according to manufacturer’s instructions and transferred to 50 mL of Human hepatocyte thawing medium (LifeNet Health). Cells were pelleted at 150 *g* for 5 minutes, the pellet was gently resuspended in 25 mL of Human hepatocyte plating medium and supplement (LifeNet Health) and pelleted again at 100 *g* for 5 minutes. Cells were resuspended again in seeding medium, and 80,000 cells/well were plated in 100 μL of medium into 96-well plates (Corning BioCoat, Collagen I coated). Before transfection, seeding medium was exchanged for human hepatocyte culture medium and supplement (LifeNet Health).

### Plasmid transfection

One day prior to transfections 12,500 HEK293T or 15,000 HeLa cells were seeded per well of a 96-well plate. If applied, treatment with small molecule compounds or DMSO control was initiated 1 hour before transfections and continued for 72 hours until harvest. AZD7648 was used at a final concentration of 1 μM (1iPEn), in combination with 3 μM PolQi1 (2iPEn) or in combination with 1.5 μM PolQi1 and 1.5 μM PolQi2 (2^+^iPEn). Cells were transfected with FuGENE HD (Promega) using a 6:1 Fugene-to-DNA ratio and 100 ng of total DNA per 96 well (75 ng of editor and 25 ng of pegRNA). For PE5, an additional 8.3 ng of nicking guide was added to the transfection mix, which is the same ratio of editor/pegRNA/nicking gRNA used previously for comparing editing efficiency between PEn and PE5 (Li et al., 2023).

### Cell electroporation with RNA

HEK293T, HeLa and HepG2 cells were seeded at 30% confluency 48 hours prior to transfection. K562 cells were resuspended at 3.5×10^5^ cells/mL 24 hours before transfection. The day of the electroporation cells were collected and washed twice with phosphate buffered saline (PBS) and resuspended in nucleofection buffer (SF/SE Cell Line 96-well Nucleofector™ Kit; Lonza) containing 2 μg of PEn mRNA and 100 pmol of synthetic springRNA or spaceRNA, at 1×10^4^ cells/μL. RNP complexes were assembled for 10 minutes at room temperature in 20 mM HEPES (pH 7.5) and 150 mM KCl buffer at a final volume of 3.36 μL, using PE2 synthetic springRNA/spaceRNA and a transfection enhancer (IDT) at a final concentration of 16.05 μM, 29.76 μM and 29.76 μM, respectively. HEK293T, HepG2 and K562 cells (2×10^5^ cells/condition) were electroporated using the SF Cell Line 96-well Nucleofector™ Kit (Lonza) and the CM-130, CN-114 or FF-120 pulse code (Nucleofector 4D; Lonza), respectively. HeLa cells (2×10^5^ cells/condition) were electroporated using the SE Cell Line 96-well Nucleofector™ Kit (Lonza) and the EH-100 pulse code (Nucleofector 4D; Lonza). Post-electroporation, cells were incubated at room temperature for 10 minutes without any disturbance and were thence transferred into pre-warmed medium in 96- (K562) or 24- (HEK293T, HeLa and HepG2) well plates and incubated at 37°C with 5% CO_2_ for 72 hours before cell collection and DNA extraction.

Primary human hepatocytes were transfected 1 day after thawing using Lipofectamine MessengerMAX Transfection Reagent (Invitrogen) using the following protocol: Lipofectamine solution was prepared by mixing 0.3 μL of Lipofectamine MessengerMAX with 4.7 μL of optiMEM (Gibco) and incubated at room temperature for 10 minutes. Meanwhile, RNA solution was prepared by diluting 170 ng of PEn mRNA and 60 ng of pegRNA with optiMEM to a final volume of 5 μL. Both solutions were combined, incubated at room temperature for 15 min and added to cells. Cells were harvested for targeted amplicon sequencing analysis after 48 hours.

### Mouse embryo electroporation

RNPs were formed at room temperature for 10 minutes in 10 μL of 2x Cas9 buffer (200 mM KCl, 40 mM HEPES) containing 12 μM PEn protein and 13 μM sgRNA. The final electroporation mix (20 μL) was prepared by combining 10 μL RNP mix with 10 μL of optiMEM and pooled zona-intact embryos (Janvier labs). Electroporation was performed using BIORAD Gene Pulser Xcell Electroporation system with the following parameters: square wave protocol, voltage: 30 V, pulse length: 3 ms, number of pulses: 10, pulse interval 100 ms, cuvette length: 1 mm. The embryos were cultivated in 24-well cell culture plates for 5 days. For lysis, single embryos were placed in 200 μL PCR strips and lysed using 10 μL 1X Modified Gitschier buffer (0.2 M Tris pH 8.8, 0.1 M (NH4)_2_SO_4_, 50mM MgCl_2_, 1.7 μM SDS, and 0.1 mg/mL Proteinase K) at 37 °C for 1 hour followed by 10 minutes heat inactivation at 85 °C. Four μL of lysates were used as PCR template to perform amplicon sequencing.

### Genomic DNA extraction and amplicon sequencing

Cells were harvested using Quick Extract solution (Lucigen). Amplicons were generated with Phusion Flash High-Fidelity 2x Mastermix (F548, Thermo Scientific) or Q5 Hot Start High-Fidelity 2x Mastermix (M0492, NEB) in a 15 μL reaction, containing 1.5-2 μL of genomic DNA extract and 0.2 μM of target-specific primers with barcodes and adapters for next generation sequencing (NGS). All primer sequences are listed in Supplementary Data 1. PCR cycling conditions for Phusion Flash High-Fidelity 2x Mastermix were: 98 °C for 3 min, followed by 30 cycles of 98 °C for 10 seconds, 60 °C for 20 seconds, and 72 °C for 30 seconds. For Q5 Hot Start High-Fidelity 2x Mastermix the following PCR protocol was applied: 98 °C for 30 seconds, followed by 30 cycles of 98 °C for 10 seconds, 60 °C for 20 seconds, and 72 °C for 30 seconds, and final elongation at 72 °C for 2 min. For off-target analysis, amplicons were generated using Q5 Hot Start High-Fidelity 2x Mastermix with the following cycling conditions: 98 °C for 3 min, followed by 30 cycles of 98 °C for 10 seconds, 65 °C for 15 seconds, and 72 °C for 30 seconds, and final extension at 72 °C for 2 minutes. Amplicons from mouse embryo lysates were generated using Q5 Hot Start High-Fidelity 2x Mastermix (M0492, NEB) in a 25 μL reaction, containing 4 μL of genomic DNA extract and 0.2 μM of target-specific primers with barcodes and NGS adapters. PCR program to generate amplicons from mouse embryos: 98 °C for 30 seconds, followed by 33 cycles of 98 °C for 10 seconds, 64 °C for 15 seconds, and 72 °C for 15 seconds, and final elongation at 72 °C for 2 minutes. All amplicons were purified using HighPrep PCR Clean-up System (MagBio Genomics). The size, purity, and concentration of amplicons were determined using a fragment analyzer (Agilent). To add Illumina indexes to the amplicons, samples were subjected to a second round of PCR. Indexing PCR was performed using KAPA HiFi HotStart Ready Mix (Roche), 0.067 ng of PCR template and 0.5 μM of indexed primers in the total reaction volume of 25 μL. PCR cycling conditions were 72 °C for 3 minutes, 98 °C for 30 seconds, followed by 10 cycles of 98 °C for 10 seconds, 63 °C for 30 seconds, and 72 °C for 3 min, with a final extension at 72 °C for 5 minutes. Samples were purified with the HighPrep PCR Clean-up System (MagBio Genomics) and analyzed using a fragment analyzer (Agilent). Samples were quantified using a Qubit 4 Fluorometer (Life Technologies) and subjected to sequencing using Illumina NextSeq system according to the manufacturer’s instructions.

### Bioinformatic analysis

Demultiplexing of the targeted amplicon sequencing data was performed using bcl2fastq software. The fastq files were analyzed using CRISPResso2 V2.1.1 in the prime editing mode (Clement et al., 2019). Detailed parameters are listed in the **Supplementary Data 1**. Histograms in Supplementary Fig. 1 were generated using CRISPResso2.

### Statistics

Data visualization and statistical analysis were conducted using GraphPad Prism 9 (GraphPad Software, Inc.) or JMP 14.1.0 (SAS Institute Inc.). Figure legends contain information on statistical tests, sample sizes, and *P* values. No data were excluded from the analyses. No statistical method was used to predetermine sample size. The experiments were not randomized. The investigators were not blinded during experiments and outcome assessment.

## Acknowledgements

We thank the AstraZeneca Discovery Sciences Genome Engineering team for support and input on this work. We thank Steve Rees and Mike Snowden for supporting this project. We are grateful to the AstraZeneca NGS & Transcriptomic team for support with targeted amplicon sequencing. P.A. is funded by the European Union’s Horizon Europe research and innovation programme under grant agreement No 101057659 (EDITSCD). L.D. is funded by Promega Corporation.

## Author contributions

P.A., L.D., N.A., M.M. and M.P. conceptualized the study and prepared the manuscript. P.A., L.D., N.A. and M.P. performed most of the experimental work and analysis with support from N.S., K.M.B., A.L.L., E.G., G.T., P.P.H., S.W., S.Š. and M.F.

## Supplementary figures

**Supplementary Figure 1:**
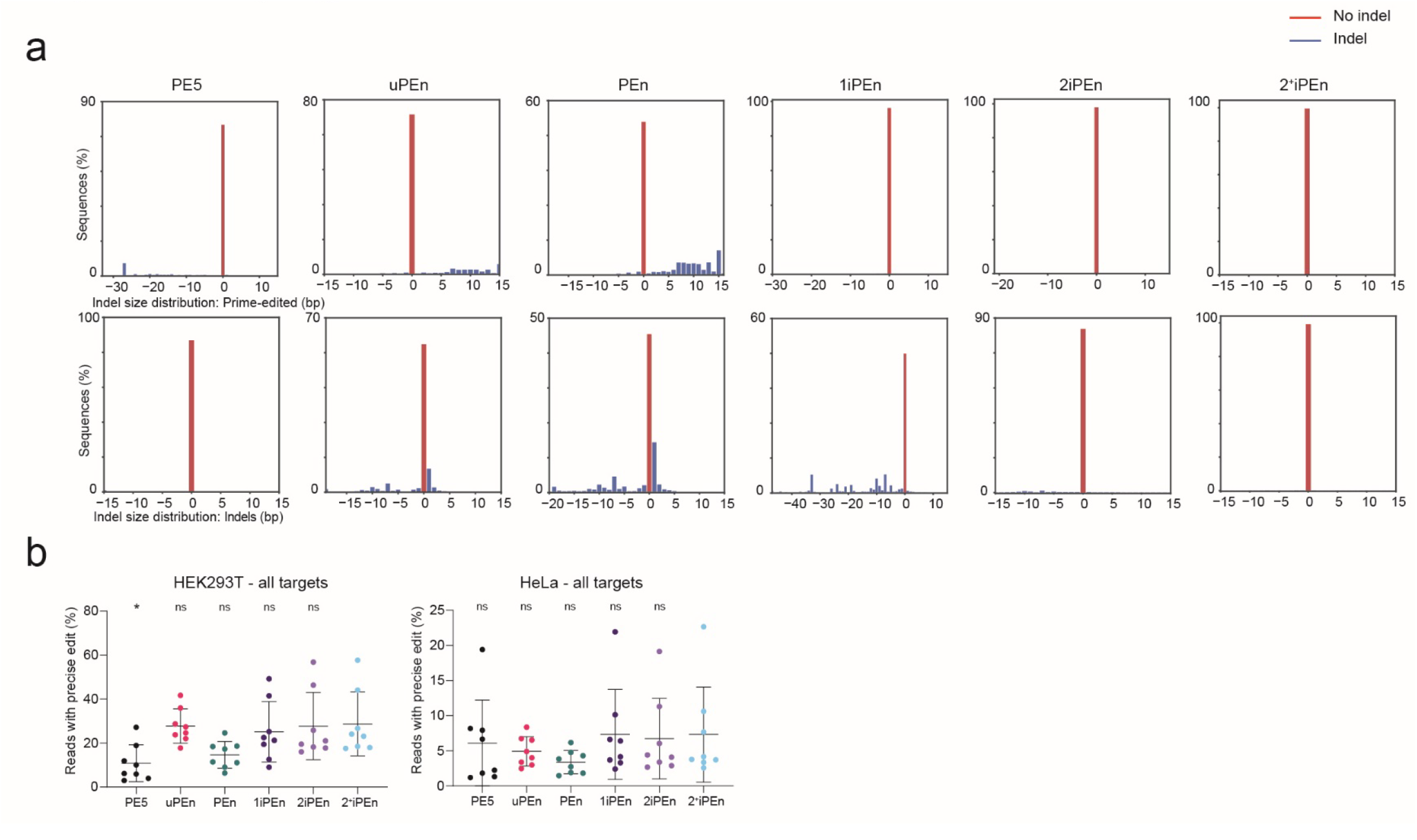
Co-inhibition of DNA-PK and PolΘ increases precision of PEn editing. **a)** Representative histograms showing size distribution of prime edits (top row) or indels (bottom row) for indicated prime editing systems. The example shows editing at the *PCSK9* locus in HEK293T cells. **b)** Prime editing efficiency of six different prime editing systems in HEK293T and HeLa cells across all loci tested in Figure 1a and b. Error bars represent mean ± SD. *P* values were determined using one-way ANOVA (two-tailed, paired). Displayed pairwise comparisons are relative to 2^+^iPEn. Calculated *P* values HEK293T cells 2^+^iPEn vs. PE5 = 0.172; 2^+^iPEn vs uPEn = 0.9996; 2^+^iPEn vs. PEn = 0.0956; 2^+^iPEn vs. 1iPEn = 0.4121; 2^+^iPEn vs. 2iPEn = 0.7793. Calculated *P* values Hela cells 2^+^iPEn vs. PE5 = 0.9994; 2^+^iPEn vs uPEn = 0.8641; 2^+^iPEn vs. PEn = 0.5253; 2^+^iPEn vs. 1iPEn >0.9999; 2^+^iPEn vs. 2iPEn > 0.8494. *P* values for all other comparisons are provided as Supplementary Data 1.

**Supplementary Figure 2:**
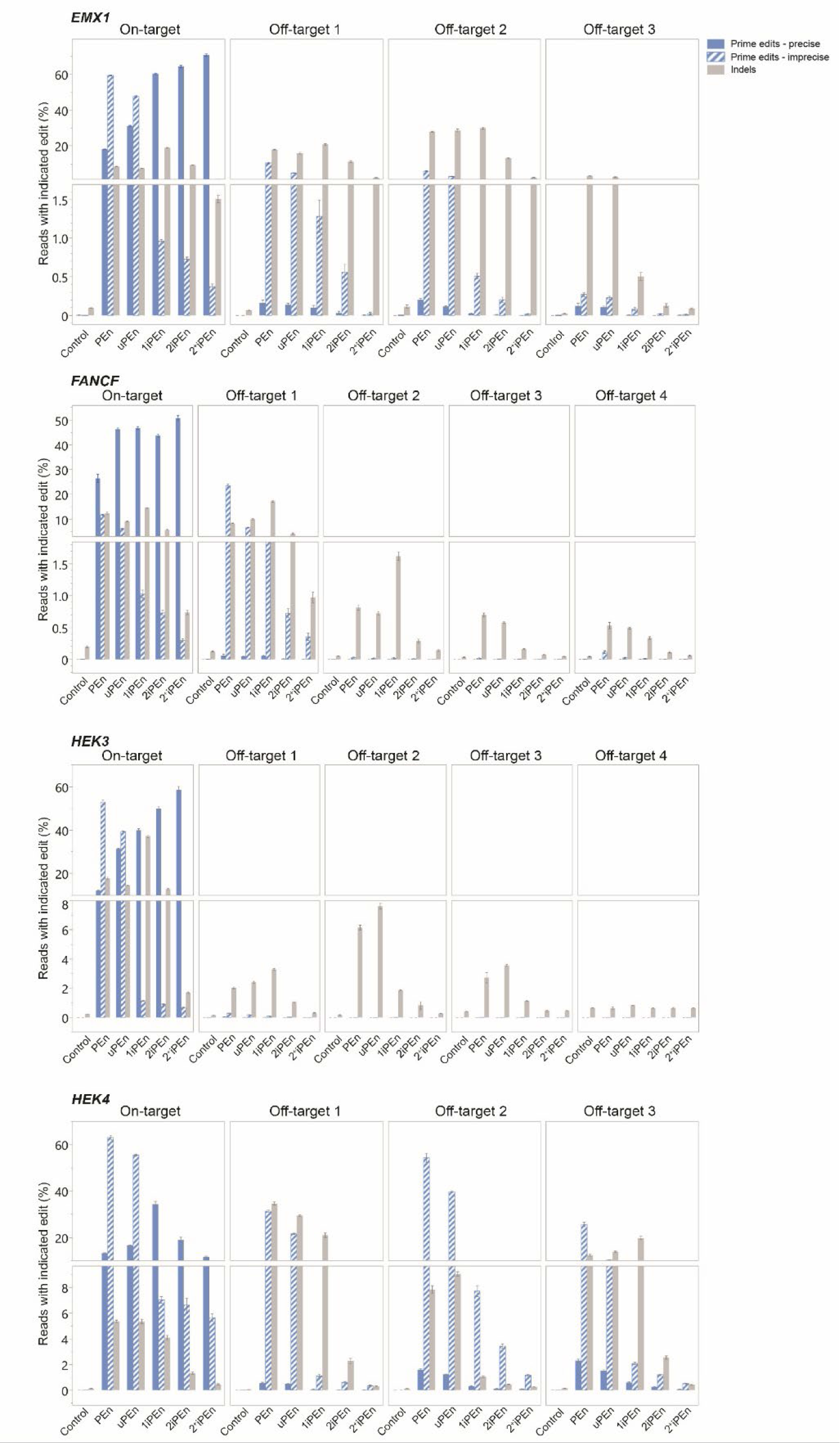
Co-inhibition of DNA-PK and PolΘ decreases PEn off-target editing. Targeted amplicon sequencing and CRISPResso 2 analysis of editing outcomes at four on-target and 13 off-target sites for the indicated PEn systems. Bar graphs represent mean values ± SEM of *n* = 3 biological replicates.

**Supplementary Figure 4:**
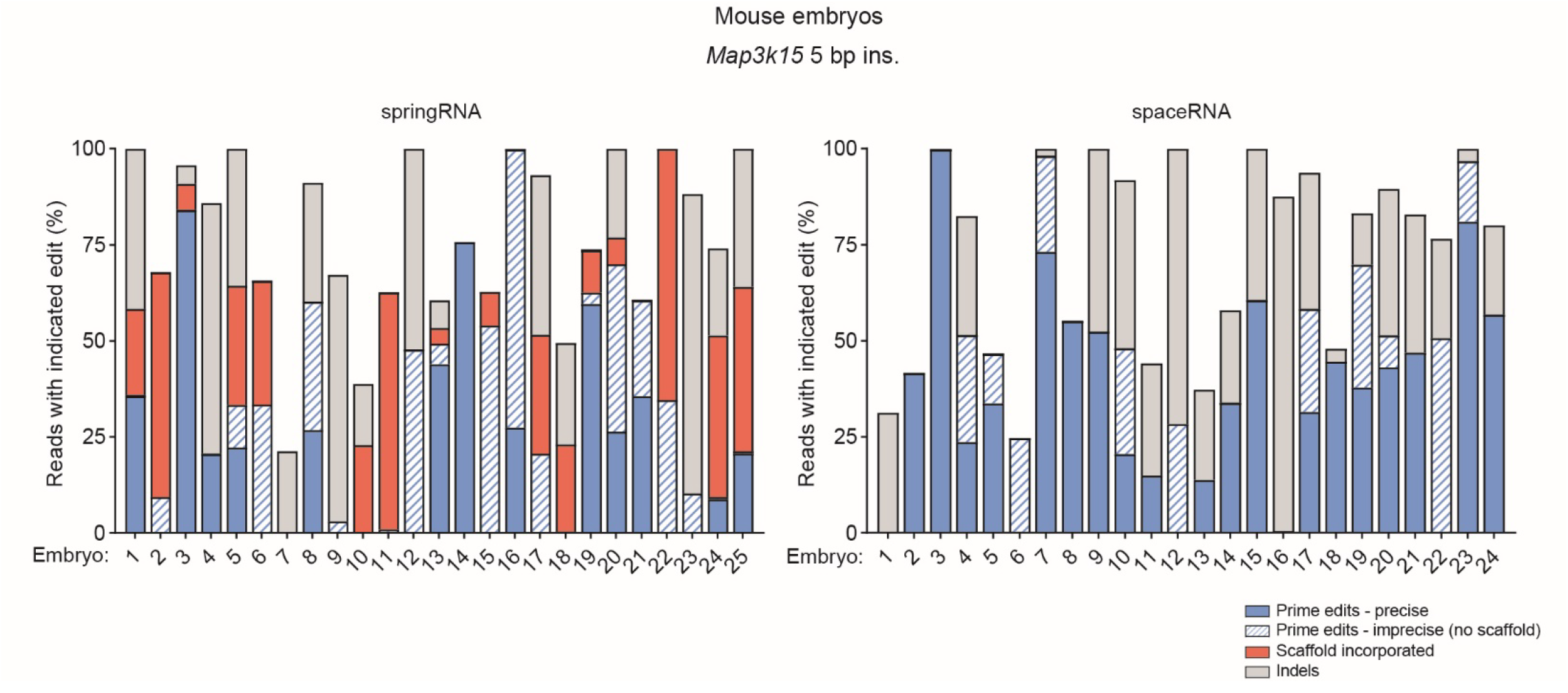
**a)** PRINS editing of one-cell mouse embryos using electroporation of PEn-springRNA or PEn-spaceRNA RNP complexes to install a 5 bp insertion into the endogenous *Map3k15* locus. Stacked bars represent indicated editing products in single embryos edited with either springRNA (left) or spaceRNA (right). Embryos were analyzed after 5 days *in vitro* cultivation by targeted amplicon sequencing and the editing outcomes were quantified using CRISPResso2.

## References

Adikusuma, F., Lushington, C., Arudkumar, J., Godahewa Gelshan I., Chey, Y. C. J., Gierus, L., Piltz, S., Geiger, A., Jain, Y., Reti, D., Wilson, L. O. W., Bauer Denis C. & Thomas Paul Q. 2021. Optimized nickase-and nuclease-based prime editing in human and mouse cells. Nucleic Acids Research, 49, 10785–10795.

Anzalone, A. V., Randolph, P. B., Davis, J. R., Sousa, A. A., Koblan, L. W., Levy, J. M., Chen, P. J., Wilson, C., Newby, G. A., Raguram, A. & Liu, D. R. 2019. Search-and-replace genome editing without double-strand breaks or donor DNA. Nature, 576, 149–157.

Arai, D. & Nakao, Y. 2021. Efficient biallelic knock-in in mouse embryonic stem cells by in vivo-linearization of donor and transient inhibition of DNA polymerase θ/DNA-PK. Sci Rep, 11, 18132.

Boiteux, S. & Guillet, M. 2004. Abasic sites in DNA: repair and biological consequences in Saccharomyces cerevisiae. DNA Repair, 3, 1–12.

Chen, P. J., Hussmann, J. A., Yan, J., Knipping, F., Ravisankar, P., Chen, P.-F., Chen, C., Nelson, J. W., Newby, G. A., Sahin, M., Osborn, M. J., Weissman, J. S., Adamson, B. & Liu, D. R. 2021. Enhanced prime editing systems by manipulating cellular determinants of editing outcomes. Cell, 184, 5635–5652.e29.

Chen, P. J. & Liu, D. R. 2023. Prime editing for precise and highly versatile genome manipulation. Nat Rev Genet, 24, 161–177.

Clement, K., Rees, H., Canver, M. C., Gehrke, J. M., Farouni, R., Hsu, J. Y., Cole, M. A., Liu, D. R., Joung, J. K., Bauer, D. E. & Pinello, L. 2019. CRISPResso2 provides accurate and rapid genome editing sequence analysis. Nature Biotechnology, 37, 224–226.

Fiumara, M., Ferrari, S., Omer-Javed, A., Beretta, S., Albano, L., Canarutto, D., Varesi, A., Gaddoni, C., Brombin, C., Cugnata, F., Zonari, E., Naldini, M. M., Barcella, M., Gentner, B., Merelli, I. & Naldini, L. 2023. Genotoxic effects of base and prime editing in human hematopoietic stem cells. Nature Biotechnology.

Fok, J. H. L., Ramos-Montoya, A., Vazquez-Chantada, M., Wijnhoven, P. W. G., Follia, V., James, N., Farrington, P. M., Karmokar, A., Willis, S. E., Cairns, J., NikkilÄ, J., Beattie, D., Lamont, G. M., Finlay, M. R. V., Wilson, J., Smith, A., O’Connor, L. O., Ling, S., Fawell, S. E., O’Connor, M. J., Hollingsworth, S. J., Dean, E., Goldberg, F. W., Davies, B. R. & Cadogan, E. B. 2019. AZD7648 is a potent and selective DNA-PK inhibitor that enhances radiation, chemotherapy and olaparib activity. Nat Commun, 10, 5065.

Jiang, T., Zhang, X.-O., Weng, Z. & Xue, W. 2021. Deletion and replacement of long genomic sequences using prime editing. Nature Biotechnology.

Kim, Y. H., Kim, N., Okafor, I., Choi, S., Min, S., Lee, J., Bae, S. M., Choi, K., Choi, J., Harihar, V., Kim, Y., Kim, J. S., Kleinstiver, B. P., Lee, J. K., Ha, T. & Kim, H. H. 2023. Sniper2L is a high-fidelity Cas9 variant with high activity. Nat Chem Biol, 19, 972–980.

Koeppel, J., Weller, J., Peets, E. M., Pallaseni, A., Kuzmin, I., Raudvere, U., Peterson, H., Liberante, F. G. & Parts, L. 2023. Prediction of prime editing insertion efficiencies using sequence features and DNA repair determinants. Nature Biotechnology, 41, 1446–1456.

Kweon, J., Hwang, H. Y., Ryu, H., Jang, A. H., Kim, D. & Kim, Y. 2023. Targeted genomic translocations and inversions generated using a paired prime editing strategy. Mol Ther, 31, 249–259.

Li, X., Zhang, G., Huang, S., Liu, Y., Tang, J., Zhong, M., Wang, X., Sun, W., Yao, Y., Ji, Q., Wang, X., Liu, J., Zhu, S. & Huang, X. 2023. Development of a versatile nuclease prime editor with upgraded precision. Nat Commun, 14, 305.

Liang, S. Q., Liu, P., Ponnienselvan, K., Suresh, S., Chen, Z., Kramme, C., Chatterjee, P., Zhu, L. J., Sontheimer, E. J., Xue, W. & Wolfe, S. A. 2023. Genome-wide profiling of prime editor off-target sites in vitro and in vivo using PE-tag. Nat Methods, 20, 898–907.

Peterka, M., Akrap, N., Li, S., Wimberger, S., Hsieh, P. P., Degtev, D., Bestas, B., Barr, J., Van De Plassche, S., Mendoza-Garcia, P., Svikovic, S., Sienski, G., Firth, M. & Maresca, M. 2022. Harnessing DSB repair to promote efficient homology-dependent and -independent prime editing. Nat Commun, 13, 1240.

Petri, K., Zhang, W., Ma, J., Schmidts, A., Lee, H., Horng, J. E., Kim, D. Y., Kurt, I. C., Clement, K., Hsu, J. Y., Pinello, L., Maus, M. V., Joung, J. K. & Yeh, J. J. 2021. CRISPR prime editing with ribonucleoprotein complexes in zebrafish and primary human cells. Nat Biotechnol.

Riesenberg, S., Kanis, P., Macak, D., Wollny, D., DÜsterhÖft, D., Kowalewski, J., Helmbrecht, N., Maricic, T. & PÄÄbo, S. 2023. Efficient high-precision homology-directed repair-dependent genome editing by HDRobust. Nature Methods, 20, 1388–1399.

Schimmel, J., MuÑoz-Subirana, N., Kool, H., Van Schendel, R., Van Der Vlies, S., Kamp, J. A., De Vrij, F. M. S., Kushner, S. A., Smith, G. C. M., Boulton, S. J. & Tijsterman, M. 2023. Modulating mutational outcomes and improving precise gene editing at CRISPR-Cas9-induced breaks by chemical inhibition of end-joining pathways. Cell Rep, 42, 112019.

Singh, P., Schimenti, J. C. & Bolcun-Filas, E. 2015. A mouse geneticist’s practical guide to CRISPR applications. Genetics, 199, 1–15.

Studier, F. W. 2005. Protein production by auto-induction in high-density shaking cultures. Protein Expression and Purification, 41, 207–234.

Tao, R., Wang, Y., Hu, Y., Jiao, Y., Zhou, L., Jiang, L., Li, L., He, X., Li, M., Yu, Y., Chen, Q. & Yao, S. 2022. WT-PE: Prime editing with nuclease wild-type Cas9 enables versatile large-scale genome editing. Signal Transduct Target Ther, 7, 108.

Tsai, S.Q., Zheng, Z., Nguyen, N.T., Liebers, M., Topkar, V.V., Thapar, V., Wyvekens, N., Khayter, C., Iafrate, A.J., Le, L.P., Aryee, M.J. & Joung, J. K. 2015. GUIDE-seq enables genome-wide profiling of off-target cleavage by CRISPR-Cas nucleases. Nature Biotechnology, 33, 187–197.

Wimberger, S., Akrap, N., Firth, M., Brengdahl, J., Engberg, S., Schwinn, M. K., Slater, M. R., Lundin, A., Hsieh, P. P., Li, S., Cerboni, S., Sumner, J., Bestas, B., Schiffthaler, B., Magnusson, B., Di Castro, S., Iyer, P., Bohlooly, Y. M., Machleidt, T., Rees, S., Engkvist, O., Norris, T., Cadogan, E. B., Forment, J. V., ŠVikoviĆ, S., Akcakaya, P., Taheri-Ghahfarokhi, A. & Maresca, M. 2023. Simultaneous inhibition of DNA-PK and PolΘ improves integration efficiency and precision of genome editing. Nat Commun, 14, 4761.

Wood, R. D. & DoubliÉ, S. 2022. Genome Protection by DNA Polymerase θ. Annual Review of Genetics, 56, 207–228.

Yeh, C. D., Richardson, C. D. & Corn, J. E. 2019. Advances in genome editing through control of DNA repair pathways. Nature Cell Biology, 21, 1468–1478.

Yu, G., Kim, H. K., Park, J., Kwak, H., Cheong, Y., Kim, D., Kim, J., Kim, J. & Kim, H. H. 2023. Prediction of efficiencies for diverse prime editing systems in multiple cell types. Cell, 186, 2256–2272.e23.

Zhao, B., Rothenberg, E., Ramsden, D. A. & Lieber, M. R. 2020. The molecular basis and disease relevance of non-homologous DNA end joining. Nature Reviews Molecular Cell Biology, 21, 765–781.

